# PRS-on-Spark (PRSoS): a novel, efficient and flexible approach for generating polygenic risk scores

**DOI:** 10.1101/209833

**Authors:** Lawrence M. Chen, Nelson Yao, Elika Garg, Yuecai Zhu, Thao T. T. Nguyen, Irina Pokhvisneva, Shantala A. Hari Dass, Marie Forest, Lisa M. McEwen, Julia L. MacIsaac, Michael S. Kobor, Celia Greenwood, Patricia P. Silveira, Michael J. Meaney, Kieran J. O’Donnell

## Abstract

**Motivation:** Polygenic risk scores describe the genomic contribution to complex phenotypes and consistently account for a larger proportion of the variance than single nucleotide polymorphisms alone. However, there is little consensus on the optimal data input for generating polygenic risk scores and existing approaches largely preclude the use of imputed posterior probabilities and strand-ambiguous SNPs.

**Results:** We developed PRS-on-Spark (PRSoS) a polygenic risk score software implemented in Apache Spark and Python that accommodates a variety of data input (e.g., observed genotypes, imputed genotypes, or imputed posterior probabilities) and strand-ambiguous SNPs. We show that PRSoS is flexible and efficient and computes polygenic risk scores at a range of p-value thresholds more quickly than existing software (PRSice). We also show that the use of imputed posterior probabilities and the inclusion of strand-ambiguous SNPs increases the proportion of variance explained by polygenic risk scores for major depression.

**Availability and Implementation:** PRSoS is written in Apache Spark and Python and is freely available (see https://github.com/MeaneyLab/PRSoS).

## INTRODUCTION

Polygenic risk scores (PRS) provide an index of the cumulative contribution of common variants to complex traits, including major depressive disorder (Peyrot et al., 2014). PRS build on large existing discovery genome-wide association studies (GWAS), such as those provided by the Psychiatric Genomics Consortium (PGC: http://www.med.unc.edu/pgc), which provide weights (odds ratios for binary outcomes and beta coefficients for continuous traits) for SNPs contributing to the PRS. PRS are given by:

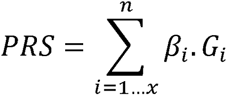

Where β_i_ = the natural logarithm of the odds ratio (or beta coefficient) between the “i^th^” SNP and phenotype of interest and G_i_ = a allele count (e.g., 0,1,2) at the “i^th^” SNP.

PRSice (Euesden et al., 2015) is written in R and uses PLINK2 (www.cog-genomics.org/plink/2.0) to calculate PRS. PRSice uses observed genotypes or posterior probabilities that have been converted to best guess genotypes (e.g., hard calls) to calculate PRS. It can also accommodate genotype dosage data but relies on a dated and slower version of PLINK (1.07) (Purcell et al., 2007). Likewise PRSice does not consider strand-ambiguous SNPs, i.e., bi-allelic SNPs that can have A-T or C-G allelic pairs, because their reference allele in the GWAS results is generally unclear. We developed a new software package, PRS-on-Spark (PRSoS), implemented in Apache Spark 2.0.0+ (Spark) and Python 2.7 that outperforms PRSice when using large genotype dosage data typical of studies in population genetics. PRSoS also provides enhanced data output options and can be applied to observed genotypes, hard calls, and imputed posterior probabilities. It further provides a reliable option to retain much of the strand-ambiguous SNPs.

### Overview of PRSoS

PRS-on-Spark (PRSoS: https://github.com/MeaneyLab/PRSoS) is implemented in Apache Spark (Spark). Spark is an open source cluster-computing framework for big data processing that can be integrated into Python programming. For the current analyses we ran PRSoS on Linux CentOS 7, 24-core Intel Xeon server with 256GB RAM, using Spark standalone mode and a distributed file system (Apache Hadoop) with 12 cores across one worker (maximum available RAM = 48GB). PRSoS can also be implemented as a stand-alone version on a single cluster. PRSoS runs on the command line in Terminal on Linux or Mac, or Command Prompt in Windows. PRSoS implements a minor allele frequency function to identify strand alignment of strand-ambiguous SNPs. PRSoS excludes strand-ambiguous SNPs that cannot be accurately aligned, e.g., SNPs with MAF >0.4 (see Supplementary Methods). PRSoS also provides an optional SNP log that documents which SNPs are included in the PRS at any given p-value threshold.

## METHODS

### Cohort

We used the PsychArray (Illumina) to describe genetic variation in women (n = 264; mean age: 35 years) from the Maternal Adversity, Vulnerability and Neurodevelopment (MAVAN) cohort (O'Donnell et al., 2014; see Table S1). We used the Center for Epidemiologic Studies Depression Scale (CES-D) to index symptoms of major depressive disorder (Radloff, 1977). See Supplementary Methods for a full description of the cohort, genotyping, quality control and imputation.

### Datasets

We compared the performance (processing times in seconds) of PRSice and PRSoS across three types of data input: imputed posterior probability/dosage data (Imputed PP), imputed genotypes converted to hard calls (Imputed HC), and observed genotypes (Array Data). PRS were calculated at five p-value thresholds (P_T_) for each of the data inputs: 0.1, 0.2, 0.3, 0.4, 0.5. However, PRSice and PRSoS are best-suited for different file formats, i.e., PLINK (.bed/.bim/.fam) format and Oxford (.gen/.sample) format, respectively. Therefore, we first compared them using the same format (Oxford files) for Imputed PP. Thereafter, we compared them using their optimal formats for all three data inputs. We calculated PRS three times for PRSoS and PRSice, and used a Student’s t-test to describe differences in total processing time. We also tested if the enhanced data output options available in PRSoS (i.e., SNP log; see Table S3 for example data output) increases PRS computation time. We also compared the performance of PRSice and PRSoS using the Imputed HC dataset to generate PRS at variable numbers of P_T_ between 0 and 0.5 to see how the number of P_T_ applied in a single run can affect processing time. We compared seven runs where PRS were generated at 5, 10, 25, 50, 100, 125, or 200 P_T_ (p-value range: 0-0.5).

## RESULTS

PRSoS calculated PRS using imputed posterior probabilities (.sample/.gen files, total input file size: 29.0GB) in 169.6sec (SD = .93sec). The same calculation using PRSice took 8461.3sec (SD = 334.6sec) (t = 42.865, p = 5.4E04; Figure 1a). Figure 1a also shows the performance of PRSice and PRSoS using the Imputed Hard Call (Imputed HC) and Array Data. PRSoS calculated PRS more quickly than PRSice when using the Imputed HC dataset (t = 62.627, p = 2.55E-04) but not when using the smaller Array Data (t = -24.978, p = 1.60E-03), where PRSice performed best. The addition of our enhanced data output (“--snp_log”) did not significantly increase processing times (see Figure S1).

**Figure 1.**
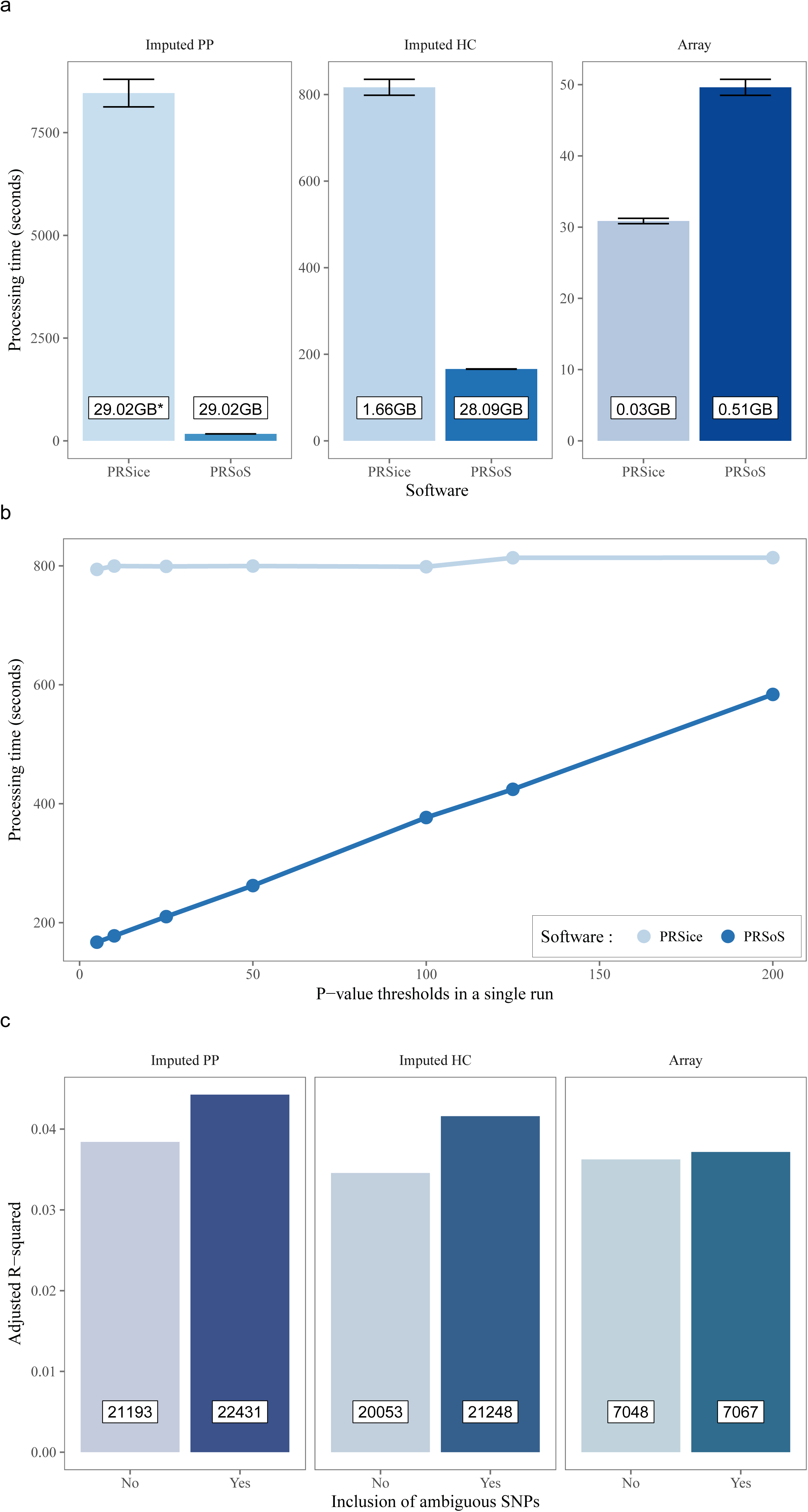
PRSoS performance. The performances of PRSoS and PRSice were compared across the dataset using imputed posterior probabilities (Imputed PP), posterior probabilities converted to hard calls (Imputed HC) and directly observed genotypes (Array) (a), and across an increasing number of p-value thresholds (b). Polygenic risk scores calculated using PRSoS based on the Imputed PP dataset that included strand-ambiguous SNPs accounted for the largest proportion of variance in symptoms of major depression (c). Numbers in inserts refer to the number of SNPs included in each polygenic risk score.

### The number of p-value thresholds affects PRSoS performance

PRSice provides a high-resolution option, creating PRS at a large number of p-value thresholds. We tested the performance of PRSoS at a range of p-value thresholds (up to 200). PRSoS processing times increase linearly with the number of p-value thresholds but PRSoS outperformed PRSice for all comparisons up to 200 p-value thresholds (see Figure 1b).

### Strand-ambiguous SNPs explain additional variance in phenotype

We tested which data input (i.e., Array Data, Imputed HC, and Imputed PP datasets with and without strand-ambiguous SNPs) provided the most informative PRS in the prediction of symptoms of major depression. We observed a positive association between PRS for MDD and maternal depressive symptoms across all datasets (see Figure S2), however the optimal P_T_ varied across different datasets. For example the PRS at P_T_ = 0.2 accounted for the largest proportion of variance of all PRS generated the Array Data. In contrast the PRS at P_T_ = 0.1 performed best for both the Imputed HC and Imputed PP datasets. PRS generated using the Imputed PP dataset that included strand-ambiguous SNPs accounts for the largest proportion of variance in depressive symptoms (adjusted R^2^ = 0.044, F(1,234) = 11.88, p = 6.7E-04). In all models, inclusion of strand-ambiguous SNPs in the PRS increases the proportion of variance that could be explained by PRS (Figure 1c).

## DISCUSSION

PRS-on-Spark (PRSoS) is a flexible and efficient software for generating PRS. We show that PRSoS, which makes use of parallel computing, outperforms PRSice when using imputed allele dosage data at a range of p-value thresholds. We also show that PRSoS accommodates strand-ambiguous SNPs, which increase the proportion of variance in depressive symptoms explained by a PRS for MDD. Future work will determine if the inclusion of strand-ambiguous SNPs increases the proportion of variance explained in other complex phenotypes.

PRSice outperformed PRSoS when using the smaller dataset of observed genotypes (Array Data), but this finding likely reflects the overhead required by PRSoS to parallelize the computation of PRS. Finally, we note that PRSice shows consistent performance across all p-value thresholds and is likely to outperform PRSoS when generating PRS at high resolution (e.g., >200 p-value thresholds).

## CONCLUSION

We have developed new software that makes use of parallel computing to accelerate PRS calculation. The increased computational efficiency of PRSoS, with its enhanced data output, will facilitate the application of PRS to better understand the polygenic basis of complex traits.

